# Maize brace roots provide stalk anchorage

**DOI:** 10.1101/2020.07.28.225656

**Authors:** Jonathan W. Reneau, Rajdeep S. Khangura, Adam Stager, Lindsay Erndwein, Teclemariam Weldekidan, Douglas D. Cook, Brian P. Dilkes, Erin E. Sparks

## Abstract

Mechanical failure, known as lodging, negatively impacts yield and grain quality in crops. Limiting crop loss from lodging requires an understanding of the plant traits that contribute to lodging-resistance. In maize, specialized aerial brace roots are reported to reduce root lodging. However, their direct contribution to plant biomechanics has not been measured. In this manuscript, we find that brace roots establish a rigid base (i.e. stalk anchorage) to limit plant deflection in maize. The more brace root whorls that contact the soil, the greater the contribution of brace roots to anchorage. Previous studies have linked the number of brace root whorls to flowering time in maize. To determine if flowering time selection alters the brace root contribution to anchorage, a subset of the Hallauer’s Tusón tropical population was analyzed. Despite a significant change in flowering time, selection neither altered the number of brace root whorls in the soil nor the overall contribution of brace roots to anchorage. These results demonstrate that brace roots provide a rigid base in maize, but the contribution to anchorage is not linearly related to flowering time.

## Introduction

### Plant Biomechanics and Lodging-Resistance

The failure of crop plants to maintain a vertical position is referred to as lodging, and can affect both crop yield and grain quality [Berry et al., 2004, Rajkumara, 2008, Fedenko et al., 2015]. Plant lodging occurs due to weak stalk and/or failed root anchorage, and multiple factors contribute to lodging susceptibility. Susceptibility factors include field management practices, meteorological factors, biotic stresses, and plant characteristics [Brune et al., 2017, Berry et al., 2004, Rajkumara, 2008, Stamp and Kiel, 1992]. Natural lodging is difficult to study due to the multiple and unpredictable factors that influence the susceptibility. To overcome this challenge, proxy measures have been developed to determine the plant characteristics that optimize for lodging-resistance [Erndwein et al., 2020]. These proxy measures rely on principles of biomechanics to link plant mechanical properties with lodging-susceptibility.

In maize, lodging has been shown to reduced yield by 3% to 25% [Flint-Garcia et al., 2003, Carter and Hudelson, 1988]. For stalk lodging, biomechanical measures of stalk breaking strength and stalk flexural stiffness have shown high association with lodging-resistance [Robertson et al., 2016, Sekhon et al., 2020]. These biomechanical properties have been further associated with variation in plant characteristics (e.g. stem morphology), and the underlying stem material properties (e.g. bending strength) [Stubbs et al., 2020, Robertson et al., 2016]. In contrast, there is a limited understanding of how root systems contribute to lodging-resistance.

### The Role of Brace Roots in Lodging-Resistance

Maize has specialized brace roots that develop in a whorl from above-ground stem nodes [Blizard and Sparks, 2020]. The uppermost whorls of brace roots may remain aerial with the lower whorls penetrating into the soil. As their name suggests, the brace roots that penetrate the soil have been proposed to provide anchorage and limit root lodging [Liu et al., 2012, Shi et al., 2019, Sharma and Carena, 2016]. Previous studies linking brace roots to root lodging-resistance have identified the brace root traits that are correlated with higher lodging-resistance as (a) a higher number of roots in a whorl [Sharma and Carena, 2016, Liu et al., 2012], (b) more overall brace root whorls entering the soil [Sharma and Carena, 2016, Shi et al., 2019], and (c) a higher brace root spread width [Sharma and Carena, 2016]. While these studies support the importance of brace roots for lodging-resistance, a direct assessment of the contribution of brace roots to plant biomechanics has not been reported.

### Genetic Regulation of Brace Roots

Although optimal brace root traits for root lodging-resistance have been identified, there are limited reports to define the genetic basis of these traits. The studies that have been reported primarily focus on the total number of brace root whorls [Zhang et al., 2018a, Ku et al., 2012, Gu et al., 2017, Zhang et al., 2018b], with only one study differentiating between the number of brace root whorls that enter the soil and the total number of brace root whorls [Ku et al., 2012]. In this study, an analysis of recombinant inbred line (ril) and immortalized F2 (if2) populations identified both shared and independent quantitative trait loci (qtl) for brace root whorls in the soil and the total number of brace root whorls [Ku et al., 2012]. One of the shared qtl on chromosome 10 explained 16.36% (ril) and 17.88% (if2) variation of total brace root whorls and 3.50% (ril) and 6.37% (if2) of the variation in the number of brace root whorls in the soil [Ku et al., 2012]. While the gene or genes underlying this major qtl have not been determined, the location overlaps with *cct1* (constans, constans-like, toc1; B73 v4: Zm00001d024909), a major determinant of photoperiod responses in maize [Hung et al., 2012]. A separate study of a maize-teosinte *BC*_2_*S*_3_ population showed significant overlap for brace root whorl number qtl and flowering time qtl [Zhang et al., 2018b], suggesting a link between the control of flowering time and brace root whorl numbers. It is unknown if modulating flowering time is sufficient to alter the number of brace root whorls in the soil, and thus affect lodging-resistance.

### Existing Methods to Quantify Lodging-Resistance

Prior studies of root lodging-resistance have employed destructive measures of pulling/pushing forces or root failure moment [Erndwein et al., 2020]. However, it is more advantageous to use non-destructive testing strategies that enable the direct measurement of brace root contribution. The Device for Assessing Resistance to Lodging IN Grains (darling) is ideal for non-destructive field-based mechanical testing [Cook et al., 2019]. Darling is a portable device equipped with a variable height load cell (force sensor) on a vertical post and a hinged foot plate equipped with an inertial measurement unit (imu). This device acquires continuous force-rotation data, which can be used to assess plant biomechanics. Previously, this device has been used to measure stalk flexural stiffness, which has been associated with stalk bending strength and stalk lodging-resistance [Robertson et al., 2016, Sekhon et al., 2020]. Measurements of stalk properties with darling assume that the plant acts as a cantilever beam with a rigid boundary condition at the base (soil). The rigidity at the base is provided by the anchorage of the root system, with both below-ground and above-ground (brace root) contributions. However, the relative contribution of these different root types has not been assessed.

In this study, we show that brace roots significantly contribute to the anchorage of maize, and that more brace root whorls in the soil leads to a greater contribution. However, modulating flowering time is not sufficient to alter the number of brace root whorls in the soil nor the relative contribution to anchorage. Together these results establish the methods for a direct assessment of the contribution of brace roots to anchorage, and highlight limitations for future studies aimed at modulating the brace root contribution to anchorage.

## Materials & Methods

### Plant material

For repeat and time of day testing, CML258 (CIMMYT tropical inbred line) seeds were planted in two replicate plots in Newark, DE during the summer of 2019. At 103 days after planting (dap) (tasseling stage - VT), 12 plants from replicate one and 11 plants from replicate two were tested for differences in the Force-Deflection slope through repeat and time of day testing. These same plants were tested 25 days later (128 dap) (reproductive dent stage - R5) and the contribution of brace roots was determined as described below. Two plants from the first replicate were unable to be tested at R5 due to damage incurred during weed control measures.

For investigating the link between flowering time and the brace root contribution to anchorage, the Hallauer’s Tusón [Teixeira et al., 2015] population was analyzed. The Tusón base population (g0) was produced by one generation of intermating between five maize accessions: PI449556, PI583912, NSL283507, PI487940, and PI498583. This base population was then subjected to open pollination in isolation and selection was made for early anthesis/silking and standability (reduced root and/or stalk lodging) for 10 generations [Teixeira et al., 2015]. Based on results from Teixeira et al. [2015], a sub-population from even-numbered generations was selected based on flowering times in Delaware closest to the mean for each generation. These lines were planted in a randomized block design in Newark, DE in the summer of 2018. Data were collected from two to three plants derived from three to five accessions in each even-numbered generation, with up to two replicated plots (Table S1). To overcome the moisture variability in green plants, samples were tested for the brace root contribution to anchorage after dry-down (154-158 dap).

### Data collection

Field-based measurements of plant biomechanics were collected with darling devices [Cook et al., 2019]. Here the devices were used to non-destructively flex the plant and extract the slope of the Force-Deflection data. The load cell was adjusted to a height of 0.64*m* or 0.60*m* from the base for CML258 and Tusón respectively, and an IMU was used to measure rotation. The difference in load cell height was unintentional, however, it is not expected to affect the results because the two datasets were not directly compared. For each test, the pivot point of the device was aligned with the base of the plant and the load cell placed in contact with the stalk. Three cycles of deflection were applied to each plant. Each cycle consisted of slowly moving the device forward to approximately 15-degrees and returning to approximately 0-degrees.

To determine if time-of-day affected test outcome when measuring green plants, two replicate plots (A and B) with 12 and 11 plants respectively were tested at 09:00 AM, 12:00 PM, and 04:00 PM. To ensure that there is no permanent deformation of the plants in response to repeat testing that would alter the response independent of brace root removal individual plants were tested three times in series. At each time point, plants one through 12 in replicate A were tested and returned to plant one to re-test each plant for a total of three repeat tests. After all three tests of replicate A were completed, replicate B was tested using the same strategy. Total time to complete the tests for one plot varied from 11-15 minutes with a total of 29-34 minutes per time point.

For testing plants during senescence or dry-down, the test with all brace roots intact was labeled “A” (Figure 1). The top whorl of brace roots that entered the soil was then manually excised with shears, and the test was repeated and labeled “B”. The process was repeated for all subsequent whorls of brace roots that entered the soil and labeled alphabetically. Here, brace roots were defined as any nodal roots visible above the soil surface.

**Figure 1:**
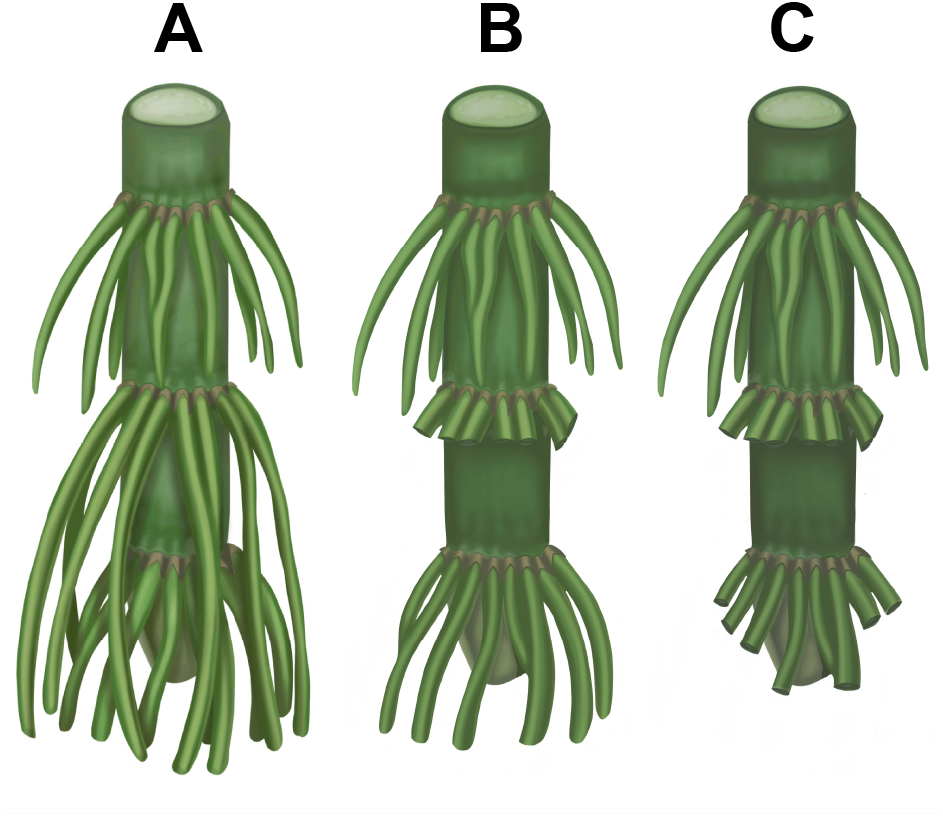
Methods for Testing the Contribution of Brace Roots to Anchorage. The measure of plant biomechanics with all brace roots intact is labeled “A”. For brace root removal experiments, the excision of sequential whorls starting at the top are labeled alphabetically. Here is an example of a plant with three whorls of brace roots, with two that enter the soil. The uppermost whorl of brace roots that enters the soil is excised and the measurement is labeled “B”. The next whorl of brace roots that enters the soil is then excised and the measurement is labeled “C”.

### Force-Deflection calculations

Cook et al. [2019] explain how darling can be used to measure the flexural stiffness of maize stalks by bending while measuring both the force and deflection. The calculation of flexural stiffness assumes that the plant acts as a cantilever beam with a rigid boundary condition at the base (soil). Under these conditions, the flexural stiffness is purely a property of the stalk itself. Here, we assume that the stalk properties are unchanging and thus any change in Force-Deflection slope from brace root removal is due changing the boundary condition of anchorage. So as not to imply that these changes are due to altering the stalk itself, here we report and analyze the Force-Deflection slope.

To convert the measured rotation to deflection, IMU data was first converted from degrees to radians with the following equation:

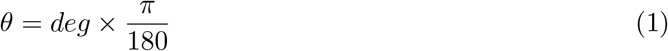

Where *π* ≈ 3.1415.

The rotation in radians was then converted into deflection (*δ*) with the following equation:

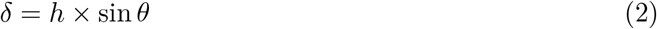

Where *h* is the height of the applied load (*m*), and *θ* is the rotational angle (*rad*) from eq. (1).

Three different approaches were compared to extract the slope from the Force-Deflection data measured from the darling devices (Figure S1A). In the first approach, a line was fit to the complete dataset, which includes loading and unloading data for all three cycles. For the second and third approaches, the data was automatically parsed into loading and unloading data points based on a user-defined number of monotonically increasing data. Here, ten consecutively increasing deflection data points were required to define a single load cycle, and each load must span at least 0.02*m* to be considered a cycle. For the second approach, a line was fit to all three loading cycles together. In the third approach, a random sample consensus (ransac) was used to fit a line to each of the cycles and the slope of the line that fit to the curve with the longest continuous data points was kept. Comparison of the different methods to extract slopes revealed that all three slopes have Pearson correlation r ≥ 0.76 and p < 2.2*E*^−16^ (Figure S1B-D). While it is ideal to extract the slope from only the loading data, the variation in the manual acquisition of data in the field environment often results in unreliable automated identification of the loading portion of the curve (e.g. wind will introduce noise that prevents the identification of monotonically increasing data points). To avoid introducing potential bias by the manual definition of loading data, the slopes of lines that were fit to the complete dataset (loading and unloading data) were used in this manuscript. Fitting a line to the complete dataset has a high correlation with fitting data to just the loading data (r = 0.86, p < 2.2*E*^−16^), thus this selection is unlikely to influence the results.

### Brace root contribution calculations

Since the removal of brace roots does not change the properties of the stalk, any differences in the Force-Deflection slope can be attributed to a change in the boundary condition and interpreted as a change in anchorage at the base. The overall brace root contribution to anchorage was calculated through relative and absolute approaches. In the relative approach, the ratio was calculated as 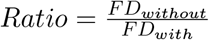, where *FD* is the Force-Deflection slope. For plants with three brace root whorls in the soil the calculation is 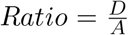 (Figure 1) and for plants with two whorls in the soil, 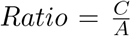. In the absolute approach, the difference was calculated as *Difference* = *FD_with_ FD_without_*, where *FD* is the Force-Deflection slope. For plants with three brace root whorls in the soil the calculation is *Difference* = *A* − *D* (Figure 1) and for plants with two whorls in the soil, *Difference* = *A* − *C*. To calculate the contribution of each whorl, the whorls were numbered beginning closest to the soil with Whorl 1 and ratios were determined as follows. For plants with three whorls, 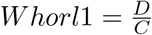, 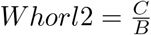, and 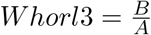. For plants with two whorls, 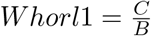, and 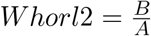.

### Genotypic Analysis of Selection

Marker genotypes for the subset of the Tusón population used in this study were obtained from [Wisser et al., 2019]. To test the effect of various plant development traits (including flowering time) on brace root anchorage across even-numbered generations of the Tusón population, we utilized phenotypes from the Newark, DE grow-out in 2009 and 2010 reported in Teixeira et al. [2015] that corresponded to the set of 26 accessions used for biomechanics measurements in this study.

### Statistics

To determine the sample sizes necessary to identify differences in the Force-Deflection slope, G*Power was used to compute sample size A priori for each of the three time points of CML258. This resulted in a sample size estimate of four. All other statistics and graphing were performed using R version 3.6.3 or JMP version 14. Specifically, one-way and two-way anova, pairwise comparisons with Tukey hsd post hoc tests and Pearson correlation analyses were performed with default parameters in R. Distributions were tested with the Shapiro-Wilk normality test, and if *p* ≥ 0.05 then the Tukey’s Ladder of Powers was used to normalize with the rcompanion package of R (version 2.3.25). Repeatability was calculated using the rptR package of R (version 0.9.22) with bootstrapping (n=1000). Graphs and input files were generated using the following R packages: ggplot2 version 3.3.0, ggpubr version 0.2.5, reshape2 version 1.4.3, and dplyr version 0.8.5. Regressions were performed in JMP.

### Data Availability

All raw data, the code used to process data, and the analyzed data are available at: https://github.com/EESparksLab/Reneau_et_al_2020.

## Results

### Plant stiffness is reduced throughout the day

The biomechanics of a plant before senescence is hypothesized to vary during the course of a day due to the variable water content (high in the morning and decreasing throughout the day), which alters turgor pressure and is thus also hypothesized to alter plant biomechanics. To quantify how biomechanics is impacted by plant physiology and time of day, two replicates of CML258 inbred plants were measured with darling at the VT stage (103 days after planting) at 9:00 AM, 12:00 PM, and 4:00 PM (Figure 2). As expected, the Force-Deflection slope is highest, indicating a stiffer plant, at 9:00 AM and decreases throughout the day (Figure 2; p = 2.361E^−06^). Thus, any biomechanical testing applied to green plants must be time-matched to circumvent these time of day effects.

**Figure 2:**
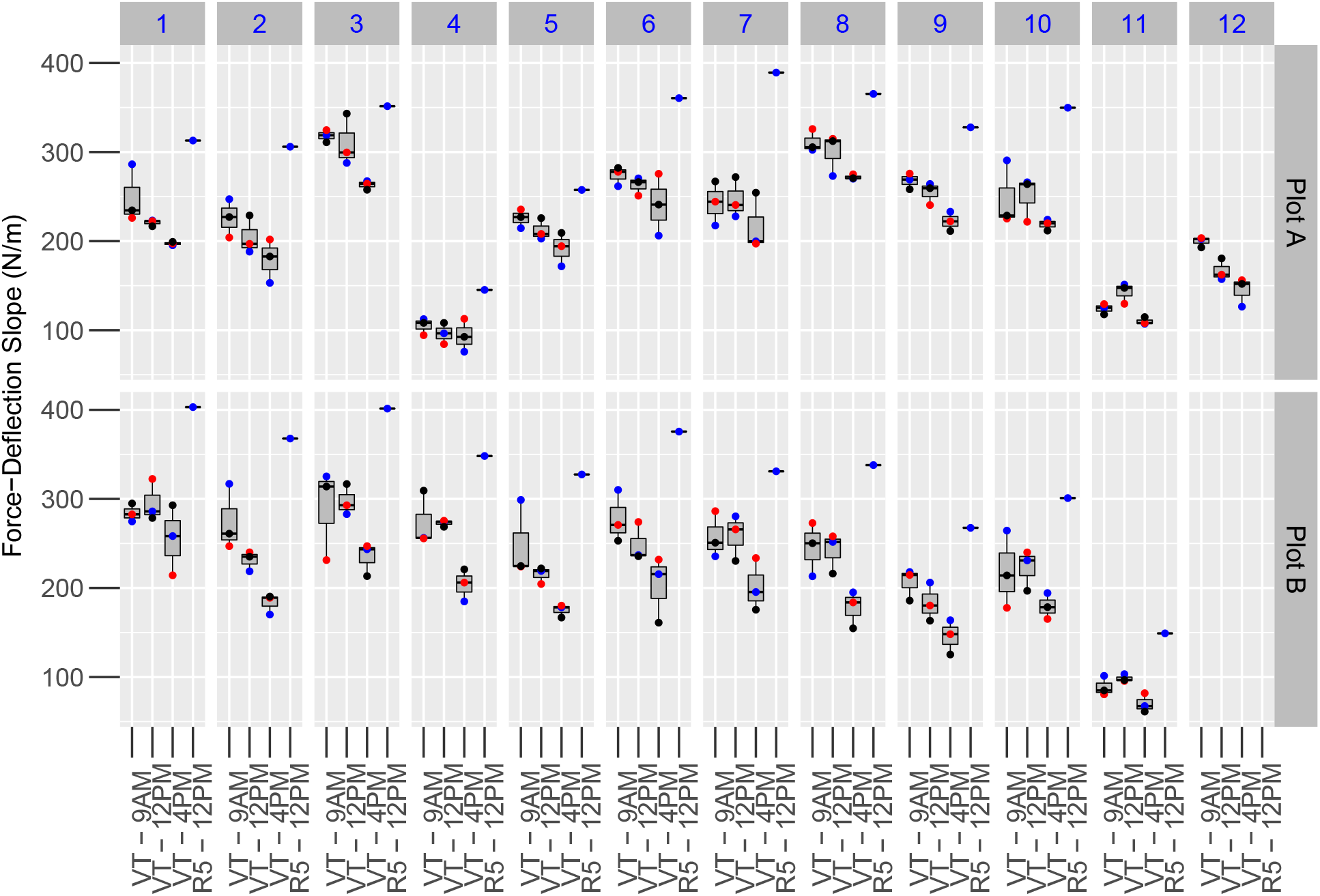
Plant Stiffness Changes with Time of Day and Developmental Stage. CML258 plants were tested at tasseling VT (103 days) and dent R5 (128 days) stages from two replicates (Plot A & B). At VT, within replicate A, each plant was tested three times as indicated by the colored dots (blue - first, red - second, and black - third tests). The Force-Deformation slope is reduced throughout the day (p = 2.36E^−06^). The effect of replication and the repeat testing was not significant (p = 0.716 and 0.972, respectively). The same plants were tested at R5 and show higher Force-Deflection slope compared to any time point at VT. Plant 11 and 12 in replicate A at R5 were not used due to weed control damage.

### Neither the number of tests nor plot replicate affects plant stiffness

Non-destructive measures of plant biomechanics are acquired under the assumption that the plant is elastically deformed and does not experience permanent deformation. To test this assumption, the same plants from replicate A and B were tested three times in a series before returning to the beginning. For each plant, there was variation in the Force-Deflection slope for the three tests, but this variation was unrelated to the test order (Figure 2; p = 0.716). Further, there was no replication effect between the two plots assayed (p = 0.972), suggesting that the time of day was the dominant factor for influencing plant biomechanics. The variation observed between repeat tests is likely due to the re-positioning of the darling foot plate relative to the plants that occurs with each subsequent test and/or the noise of data collection that impacts the extraction of the slope from the Force-Deflection data. This variation represents the technical variation that is introduced by manual field-based measurements. Despite this variation, a high repeatability is found for the Force-Deflection slope measurements (R = 0.876, SE = 0.0245, p=2.31E^−49^; Figure S2). These results indicate that neither repeat testing nor plot replication affects the Force-Deflection slope in the CML258 inbred line, and there is high repeatability with the non-destructive measurement of plant biomechanics with the darling.

### Plant stiffness is higher during senescence than the tasseling stage

To remove the impact of time of day on the Force-Deflection slope, all subsequent analyses were conducted during plant senescence and just prior to or immediately after harvest. The same CML258 plants that were tested at VT were tested again at R5 prior to harvest. There was a consistent increase in the Force-Deflection slope at R5 (Figure 2) compared to plants measured at VT. There was again no effect of plot replication on the Force-Deflection slope at R5. This data suggests that plant biomechanics change across the plant lifespan, with increasing stiffness during reproductive development relative to late vegetative development.

### Brace roots significantly contribute to plant anchorage

Brace roots are suggested to contribute to plant anchorage and root lodging-resistance, but their contribution has not been directly tested. CML258 plants have two to three whorls of brace roots that enter the soil (Figure 3A). Although there was no effect of plot replication on the Force-Deflection slope in Figure 2, the number of brace root whorls in the soil is significantly influenced by plot replication in this experiment (p = 0.0308). To determine if brace roots significantly contribute to anchorage, plants were tested after excision of successive whorls of brace roots starting from the highest whorl (Figure 1). The repeat testing of the same plant does not introduce significant variation in the Force-Deflection slope (Figure 2), so any effect can be attributed to the absence of brace roots.

**Figure 3:**
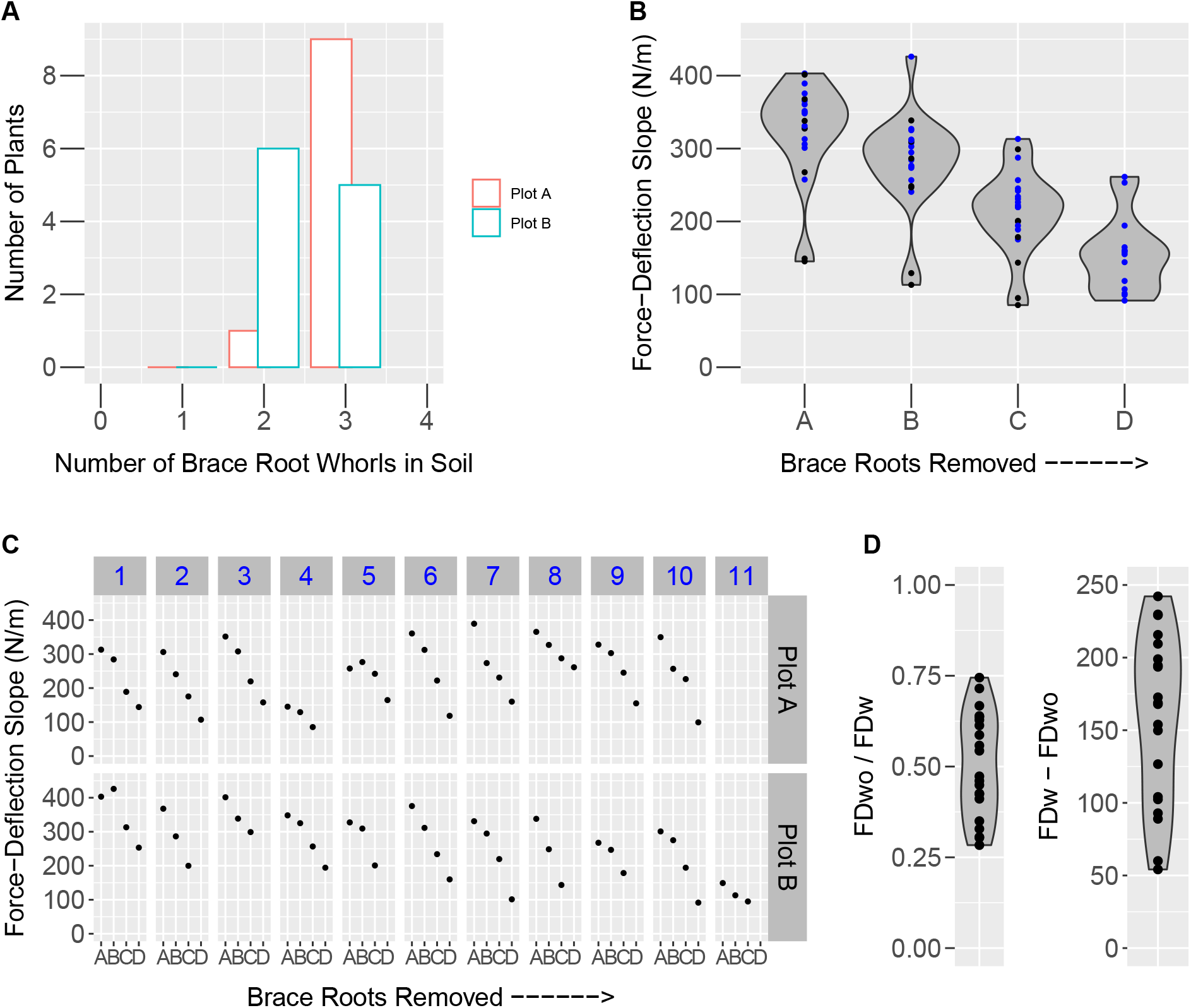
Brace Roots Contribute to Anchorage. (A) The number of brace root whorls in the soil was evaluated for CML258 plants in two replicates and there was a significant effect of replicate (p = 0.0308). (B)The Force-Deformation slope was measured from CML258 plants upon sequential removal of brace root whorls. Test was performed with all whorls intact A, excision of the highest whorl B, the next highest whorl C and so on. Plants with two whorls are represented by black and with three whorls by blue dots. There is an overall reduction in the slope upon removal of brace root whorls (p = 5.88E^−11^). (C) This observation is extended to the individual plant, where the subsequent removal of brace root whorls results in a reduction of the Force-Deflection slope. (D) Using the paired data measurements, the contribution of brace roots to anchorage is represented in two ways: 1) a ratio of the Force-Deflection slope with no brace roots to the Force-Deflection slope with all brace roots, and 2) the difference in Force-Deflection slope with all brace roots and the Force-Deflection slope with no brace roots. *FD_wo_* = Force-Deflection slope without brace roots. *FD_w_* = Force-Deflection slope with brace roots.

Brace root removal has a significant effect on the Force-Deflection slope (p = 5.88E^−11^; Figure 3B). Removal of the top whorl of brace roots does not result in a significant change to the slope (Tukey hsd A vs. B, p = 0.138), however, there is a significant reduction with removal of subsequent whorls (Tukey hsd B vs. C, p = 0.004; C vs. D, p = 0.048). The top whorl was often composed of only a few brace roots with limited penetration into the soil (*data not shown*) and thus the low impact of this whorl is expected. This collective observation is maintained at the individual plant level, with each plant showing a reduction in Force-Deflection slope upon successive removal of brace root whorls (Figure 3C). For plot A plant 5 and plot B plant 1, the B measurement is higher than the A measurement (Figure 3C), which is again consistent with the top whorl of brace roots contributing very little. The difference between these two measurements can be attributed to the same technical variation we identified in Figure 2.

Given the variation in both the number of brace root whorls (Figure 3A), and initial Force-Deflection slope (Figure 2), individual paired analyses for each plant can provide quantitative information on the overall contribution of brace roots to anchorage. The contribution of brace roots was calculated as both 1) relative as a ratio of Force-Deflection slope with all brace roots excised to the initial Force-Deflection slope and 2) absolute as a difference between initial Force-Deflection slope and Force-Deflection slope with all brace roots removed (Figure 3D). For either the ratio or difference measures, there was no significant effect of replication on these measures. There was a moderate correlation between the difference measure and the initial Force-Deflection slope (Figure S2A, r = 0.59, p = 0.004). This is not surprising, as the higher the initial Force-Deflection slope the more potential there is to reduce the slope upon brace root removal. However, when normalizing for the relative contribution by use of a ratio, the correlation disappears (Figure S2B, r = −0.17, p = 0.461). The initial Force-Deflection slope is predominantly determined by the stalk properties, and thus we would not expect any correlation with the relative brace root contribution to anchorage. To separate from the initial stalk mechanics, we utilize the relative ratio measurements for the remainder of this study. These ratios are a quantification of the relative contribution of brace roots to anchorage, and a lower value indicates a higher contribution from the brace roots.

### Brace root whorls have differential contribution to anchorage

Despite a consistent reduction in Force-Deformation slope upon removal of brace root whorls, there is variation in the relative contribution within the CML258 inbred line (Figure 3D). Previous studies have shown that the number of brace root whorls that enter the soil is correlated with root lodging-resistance [Sharma and Carena, 2016, Liu et al., 2012]. Thus, we examined the role of different number of brace root whorls in the soil.

There is a significant correlation between the brace root contribution ratio and the number of brace root whorls in the soil (Figure 4A, r = −0.50, p = 0.02). However, there is not a 50% greater brace root contribution for plants with three brace root whorls compared to plants with two brace root whorls, as would be expected if each whorl contributed equally. Analysis of the contribution of each whorl shows that the brace root whorl closest to the soil (Whorl 1) contributes the most and each additional whorl contributes successively less (Figure 4B; WR1 v. WR2 p = 0.002; WR2 v. WR3 p = 0.05). There is however a significant correlation between the contribution of all whorls and the contribution of Whorl1 (Figure 4C, r = 0.85, p = 1.302E^−06^), Whorl2 (r = 0.51, p = 0.018) or Whorl3 (r = 0.62, p = 0.017). These results suggest that, although the brace root whorl closest to the soil is contributing the most towards anchorage, the relative contribution of each whorl is directly related to the overall contribution.

**Figure 4:**
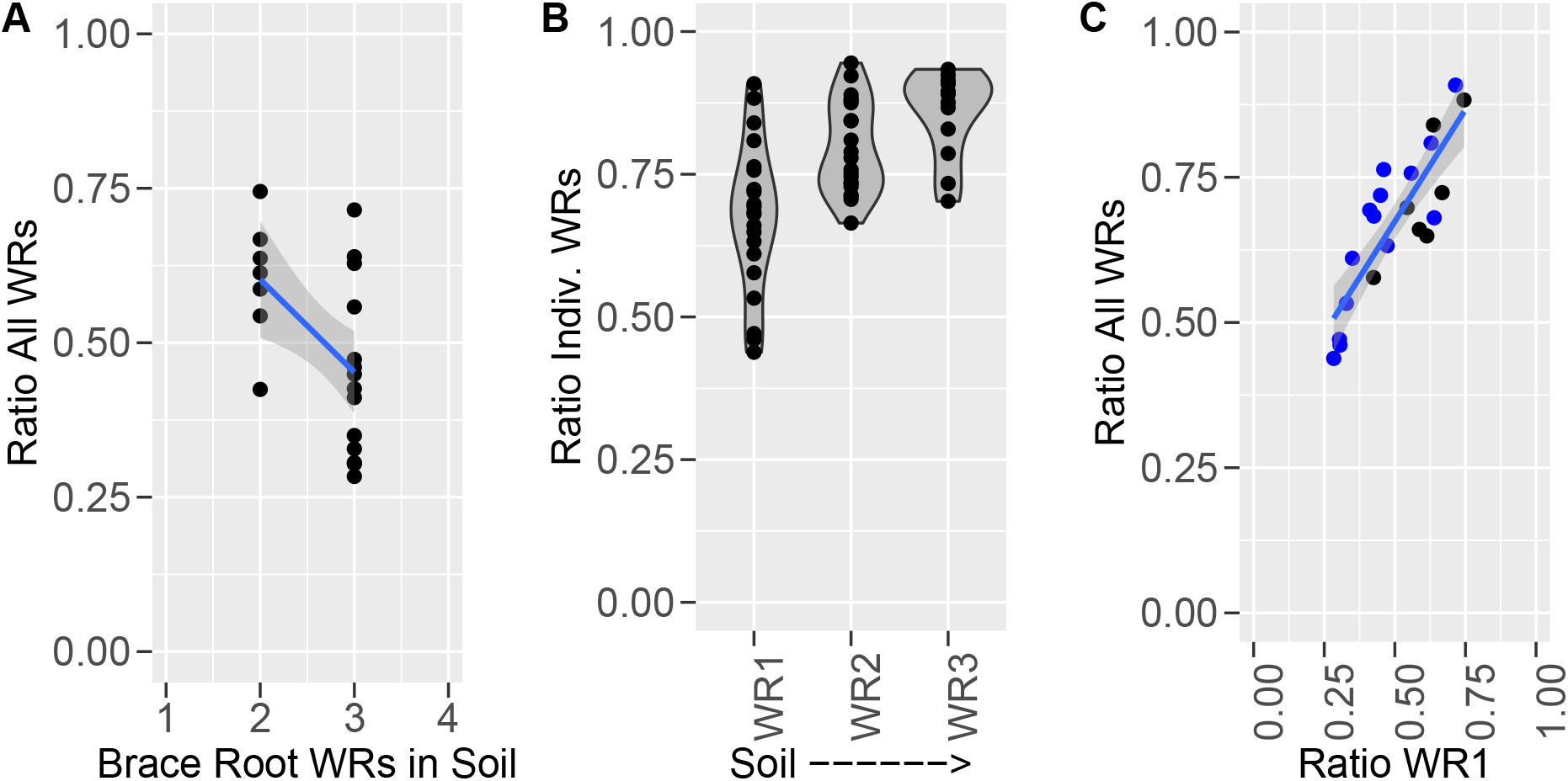
Brace Root Whorls Have Differential Contribution to Anchorage. (A) The reduction of Force-Deflection slope after removal of all brace root whorls is significantly correlated with the number of whorls in the soil (r = −0.50, p = 0.02). (B) Each whorl of brace roots in the soil has a differential contribution to the Force-Deflection slope. The effect of whorl is significant by anova, Pr(>F) = 2.4431E^−06^. The brace root whorl closest to the soil contributes the most and the additional whorls above contribute successively less. (C) The contribution of the lowest whorl is highly correlated with the overall contribution (r = 0.85, p = 1.302E^−06^). Plants with two whorls are represented by black and with three whorls by blue dots. Lines represent generalized linear model (glm) fit and shading indicates a 95% confidence interval. A lower ratio indicates a higher contribution of brace root whorls. WR = whorl.

### Selection for early flowering does not affect the number of brace root whorls in the soil or the brace root contribution to anchorage

Previous work has described a relationship between the number of nodes producing brace roots and flowering time in maize [Zhang et al., 2018b]. However, the described relationship does not take into account the number of brace root whorls that enter the soil, nor their relative contribution to anchorage. We set out to determine if selection for flowering time can alter the number of brace root whorls that enter the soil, and thus alter the contribution of brace roots to anchorage. For this analysis, a subset of lines from the Tusón population were analyzed [Teixeira et al., 2015]. The Tusón population underwent selection for early flowering time and secondary selection for standability (reduced root and stalk lodging) in central Iowa for 10 generations. The selection for early flowering in the Tusón population resulted in indirect selection for additional traits such as plant height [Teixeira et al., 2015]. A random sample of plants from even-numbered generations of the Tusón population were analyzed to determine if the selection for early flowering has indirectly selected for the number of brace root whorls in the soil, the contribution of brace roots to the Force-Deflection slope, or the initial Force-Deflection slope. Although the subset of lines analyzed showed a significant reduction in flowering time (Figure S4), the number of brace root whorls in the soil is unchanged between selection cycles (Figure 5A, p = 0.464). Consistent with this observation, the relative contribution of brace roots is also unchanged over the course of selection (Figure 5C, p = 0.115).

**Figure 5:**
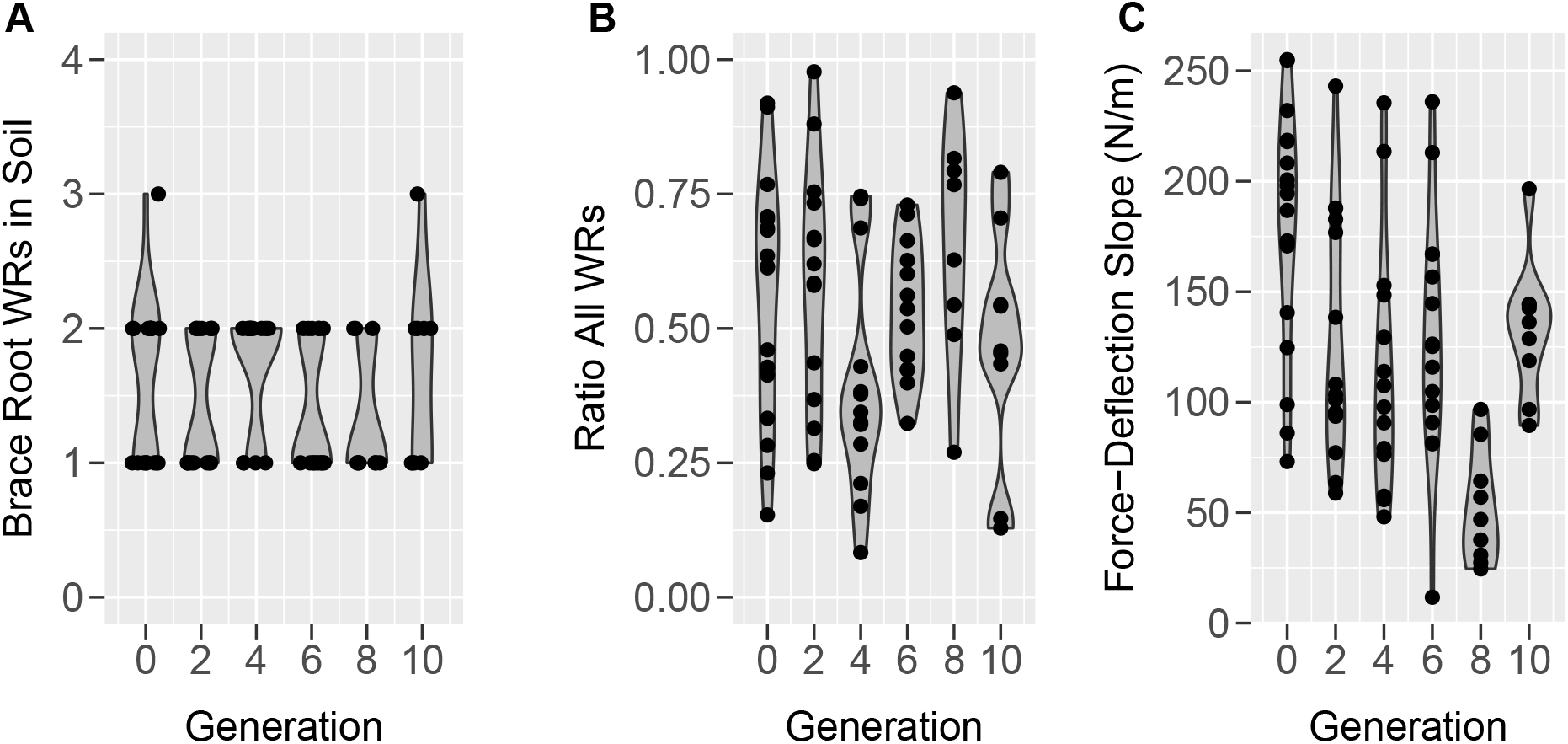
Selection for Early Flowering Alters the Force-Deflection Slope, but not the Number of Brace Roots in the Soil or the Contribution of Brace Roots to Anchorage. (A) There is no effect of flowering time selection on the number of brace root whorls in the soil. (B) The contribution of brace roots to the Force-Deflection slope is not affected through selection. (C) Selection for early flowering in the Tusón population has a significant effect on the Force-Deflection slope (p = 7.44E^−06^). WR = whorl.

### The effects of plant height and flowering time on Force-Deflection slope are separable

In contrast to the brace root traits, the initial Force-Deflection slope was significantly changed in this population (Figure 5C, p = 7.44E^−06^), and correlated with multiple phenotypes in single variable regressions (Table S2). However, stepwise inclusion of variables in the phenotypic regression indicated that these associations are largely due to trait correlation and not selection. Inclusion of flowering time in multiple regressions by considering the trait used for selection in the Tusón population, Days to Silking (dts), revealed additional relationships between Force Deflection slope and plant size traits. Dts was the most explanatory value for the initial Force-Deflection slope, and its inclusion in multiple regressions removed the association between the initial Force-Deflection slope and all other traits except the number of brace root whorls in the soil and plant height (ph). The best model retained dts and ph as significant predictors of Force-Deflection slope with an R^2^ of 0.40 (dts: effect estimate = 0.43, p = 0.001; ph: effect estimate = −0.06, p = 0.03). This suggests that as dts decreased, plant stiffness also decreased. In this best multiple regression model, the effect estimates were positive for dts (0.43) and negative for ph (−0.06). Thus, controlling for the effect of flowering time reveals a separate and opposing effect of ph on stiffness. The opposing effects of these variables is remarkable, considering that the dts and ph are positively correlated (r = 0.8; Table S2) with each other across the selection experiment (Teixeira et al. 2015). Similar regressions were done using plant per se data, instead of line means. The plant per se data uncovered the same relationships of dts and ph to the Force-Deflection slope (*data not shown*).

Closer inspection of the changes in the Force-Deflection slope across the generations of selection shows that significant changes did not occur until the eighth generation (g8; Figure 5C), where there was a significant reduction from g6 (Tukey hsd p = 0.009) followed by an increase in the Force-Deflection slope at g10 (Tukey hsd p = 0.015). At these same generations, a change in flowering time was not significant (Tukey hsd p = 0.83; Figure S4). Mean ear height also increased between g8 (119 cm) and g10 (126 cm), and mean ph decreased marginally between g8 (248 cm) and g10 (243 cm) for the lines used in this analysis (Figure S4). Although these differences were not statistically significant, the greater selection efficiency for lodging resistance and co-selection of ph in later generations of the Tusón population may explain the bounce observed in the Force-Deflection slope in g10 (Figure 5C). Indeed, greater selection efficiency for standability in later generations may select for specific plant architectures and explain the increase in the Force-Deflection slope in g10.

## Discussion

### Plant biomechanics is variable by time of day and growth stage

Analysis of the Force-Deflection slope over the course of a day and at two developmental stages revealed variable plant biomechanics (Figure 2). Change in the Force-Deflection slope over the course of the day (high in the morning and decreasing throughout the day) is consistent with variable water content altering the turgor pressure and thus plant biomechanics. These data also show that plants undergoing senescence have a higher Force-Deflection slope (stiffness) than the same plants at tasseling stage. There are three hypotheses that may explain these differences, 1) There may be differences in the elasticity of living cells versus dying/dead cells during senescence. 2) This difference may be attributed to a compensatory mechanical response as the floral organs develop creating a weighted beam. 3) Lastly, we cannot rule out that the higher Force-Deflection slope during senescence is a thigmotropic response to the plants being tested while green. However, the displacement of the stalk during testing is no more or less than we would expect from wind, people, or animals moving through the field. Thus, we do not expect the testing itself to induce additional changes to the biomechanics. While additional work is required to fully define the underlying biology of these dynamic changes in Force-Deflection slope, these results highlight the importance of field-based non-destructive tests to capture biomechanical dynamics.

### Brace roots provide anchorage

The measured Force-Deflection slope has a significant contribution from brace roots, which provide a rigid attachment base for the maize stalk (Figure 3). This contribution is likely due to brace roots acting as guy-wires to provide stability to the free-standing stem. In this way, the removal of brace roots will effectively lengthen the beam - in other words, move the anchorage point on the stem from the point of brace root attachment to the soil surface. One hypothesis is that the role of brace roots can be explained by this change in effective beam length. However, when accounting for the attachment height of brace roots (ranging between 0.015*m* to 0.032*m*, *data not shown*), there is still a significant effect of brace roots on the Force-Deflection slope. Therefore, the contribution of brace roots cannot be explained by changing the length of the beam, and instead demonstrates that brace roots contribute to stalk anchorage by providing a stable base.

### More brace root whorls have a greater contribution to anchorage

Our results demonstrate that the more brace root whorls that enter the soil, the greater the overall reduction in Force-Deflection slope (Figure 4). While this is again consistent with the idea of brace roots functioning as guy-wires, there are some notable differences with this analogy. With guy-wires, the higher set of wires contribute more than the lower set. With brace roots the lowest whorl contributes the most and the higher whorls successively less. We hypothesize that this is due to the depth of the roots in the soil; with the lowest whorl having more time to grow subterranean and provide anchorage. However, determining the depth of brace roots in the soil is a difficult challenge due to the complexity of the maize root system and limited field-based root phenotyping approaches [Clark et al., 2020].

### Selection for flowering time does not affect the relative contribution of brace roots to anchorage

Previous work has linked the number of brace root whorls with flowering time [Zhang et al., 2018b, Ku et al., 2012]. However, it was unknown if modulating flowering time affects the number of brace root whorls in the soil or their contribution to anchorage. Although there was a significant change in flowering time across selection in these population (Figure S4), there was no change in the number of brace root whorls in the soil nor their relative contribution to anchorage (Figure 5A-B).

The lack of a relationship between flowering time and brace root traits may result from the source of flowering time variation in this particular subset of the Tusón population. In the complete Tusón population twelve loci were identified that explain 60% of selected change in flowering time adaptation across the 10 cycles of selection [Wisser et al., 2019]. Of these, the non-photoperiod sensitive allele of a major determinant of photoperiod responses in maize, *cct1* was highly relevant for the flowering time selection [Wisser et al., 2019]. Due to the selection of “average” lines for our analysis, four of these loci (including *cct1*) were monomorphic in all generations including g0 and g2 and another three loci had a single line with an alternative allele (Table S3). Of the remaining five markers, only two have more than five lines with alternative alleles and none of these alleles were significantly associated with time to maturity, or the Force-Deflection slope as determined by regression analyses (*data not shown*). Despite the absence of variation in these major photoperiod genes, the subset of lines evaluated in this study still shows a significant decrease in days to anthesis and silking in Delaware (Figure S4A-B; p ≤ 0.0001; [Teixeira et al., 2015]). Thus there is additional flowering time allelic variation in this subset of lines that underlies the selection for flowering time that cannot be accounted for by the alleles detected by Wisser et al. [2019]. The absence of an effect of flowering time variation on brace root traits indicates that brace root-mediated anchorage can be controlled independent of flowering time variation and photoperiodism.

### There is a separable and opposing effect of plant height and flowering time on plant stiffness

Interpretations of the relationship between the initial Force-Deflection slope (stiffness) and changes in flowering time over the selection cycles in the Tusón population is complicated by a few factors. The first is the small sample size of the accessions evaluated in this study. For instance, the lines sampled as representatives for selection cycle g10 have higher mean ear height than g8. This is not the case for the Tusón population which flowers earlier and shows gradual and significant decrease (Tukey’s hsd p<0.05) in ear height from g8 to g10 [Teixeira et al., 2015]. The fortuitous selection of lines with unusual height in this subset may confuse the predicted slope for calculations of genetic effects of selection across the experiment, but may have helped to expose opposing effects of ph and dts regulators on plant stiffness.

Additionally, poor sampling of alternate alleles at the significant flowering time loci in this population may limit the detection of associations between dts and the Force-Deflection slope. Flowering time in maize is a complex trait [Buckler et al., 1993] and is regulated by numerous small effect loci especially in landraces which exhibit high allelic diversity [Romero Navarro et al., 2017]. Consistent with a polygenic control of flowering time, even in the absence of major flowering time loci, there is still sufficient variation in flowering time to detect a positive correlation with stiffness in these 26 lines. The absence of segregation at the marker positions from Wisser et al. [2019] suggest that the flowering time variation in this subset of Hallauer’s Tusón population is controlled by previously undetected loci.

Overall, these results are the first report of repeat measures to uncover the dynamics of plant biomechanics in the field. These results also provide the first direct measurement of the brace root contribution to anchorage. The number of brace root whorls in the soil is important for the overall contribution to anchorage. However, selection for flowering time alone is not sufficient to alter the number of brace roots in the soil or their contribution to anchorage. These results provide the foundation for future studies aimed at uncovering the variation in brace root contribution to anchorage, defining the genetic basis of this trait, and uncovering the brace root phenotypes (in addition to whorl number) that alter the contribution to anchorage.

## Acknowledgements

We would like to acknowledge members of the Sparks lab for reading and providing feedback on this work, and Dr. Randall Wisser for providing Tusón seed stocks. This work was supported by grants from the Delaware Biosciences Center for Advanced Technology, the University of Delaware Research Foundation, and the Thomas Jefferson Fund / FACE Foundation to EES.

## Supplemental Information

**Table S1:** Table of Tusón accessions and replication used in this study. Accessions are labeled as in [Teixeira et al., 2015]. Specifically, “C” indicates the generation number followed by row ID and plant number. Plot Rep indicates the number of plots that were assayed, and the number of plants is the total number across all plots.

| Accession | Generation | Plot Rep | Plants |
| --- | --- | --- | --- |
| Tuson:C0-164(5) | g0 | 1 | 3 |
| Tuson:C0-165(1) | g0 | 2 | 6 |
| Tuson:C0-165(2) | g0 | 1 | 3 |
| Tuson:C0-165(3) | g0 | 1 | 3 |
| Tuson:C0-169(1) | g0 | 1 | 3 |
| Tuson:C2-172(2) | g2 | 2 | 6 |
| Tuson:C2-172(3) | g2 | 1 | 3 |
| Tuson:C2-830(2) | g2 | 1 | 3 |
| Tuson:C2-832(2) | g2 | 1 | 3 |
| Tuson:C4-176(4) | g4 | 1 | 3 |
| Tuson:C4-176(5) | g4 | 1 | 3 |
| Tuson:C4-177(6) | g4 | 1 | 3 |
| Tuson:C4-178(6) | g4 | 1 | 3 |
| Tuson:C4-839(2) | g4 | 1 | 2 |
| Tuson:C6-849(4) | g6 | 1 | 3 |
| Tuson:C6-852(3) | g6 | 1 | 2 |
| Tuson:C6-856(5) | g6 | 1 | 3 |
| Tuson:C6-857(8) | g6 | 2 | 5 |
| Tuson:C6-858(4) | g6 | 1 | 2 |
| Tuson:C8-860(1) | g8 | 1 | 3 |
| Tuson:C8-861(1) | g8 | 1 | 3 |
| Tuson:C8-864(3) | g8 | 1 | 3 |
| Tuson:C10-869(6) | g10 | 1 | 3 |
| Tuson:C10-872(2) | g10 | 1 | 3 |
| Tuson:C10-873(1) | g10 | 1 | 3 |

**Table S2:** Trait correlations between Force-Deflection slope and plant development traits of the subset of 26 accessions from the Tusón population from [Teixeira et al., 2015]. Upper triangle (above the diagonal) shows the Pearson correlation coefficients (r), and lower triangle contains p-value for each pairwise correlation comparison. Gen = generation of selection; Ratio = Force-Deflection slope with no brace roots divided by Force-Deflection slope with brace roots intact; dta = days to anthesis; dts = days to silking; asi = anthesis-silking interval; ph = plant height; erh = ear-to-tassel length; peh = ear height; tsw = tassel weight; spl = tassel spike length.

|  | Gen | Force-Deflection | Ratio | DTA | DTS | ASI | PH | ERH | PEH | TSW | SPL |
| --- | --- | --- | --- | --- | --- | --- | --- | --- | --- | --- | --- |
| Gen | – | -0.423 | -0.01 | -0.94 | -0.92 | -0.67 | -0.83 | -0.89 | 0.14 | -0.22 | -0.11 |
| Force-Deflection | 0.03 | – | -0.42 | 0.49 | 0.51 | 0.41 | 0.17 | 0.31 | -0.31 | 0.37 | -0.04 |
| Ratio | 0.97 | 0.03 | – | -0.02 | -0.02 | -0.01 | -0.01 | 0.06 | -0.16 | -0.17 | -0.18 |
| DTA | <0.0001 | 0.01 | 0.91 | – | 0.96 | 0.66 | 0.83 | 0.86 | -0.08 | 0.30 | 0.005 |
| DTS | <0.0001 | 0.01 | 0.92 | <0.0001 | – | 0.84 | 0.80 | 0.83 | -0.08 | 0.38 | -0.04 |
| ASI | 0.00 | 0.04 | 0.96 | 0.00 | <0.0001 | – | 0.55 | 0.57 | -0.06 | 0.47 | -0.11 |
| PH | <0.0001 | 0.40 | 0.97 | <0.0001 | <0.0001 | 0.004 | – | 0.90 | 0.21 | 0.12 | -0.01 |
| ERH | <0.0001 | 0.12 | 0.77 | <0.0001 | <0.0001 | 0.002 | <0.0001 | – | -0.24 | 0.20 | -0.05 |
| PEH | 0.49 | 0.12 | 0.45 | 0.70 | 0.70 | 0.76 | 0.30 | 0.25 | – | -0.18 | 0.08 |
| TSW | 0.28 | 0.06 | 0.40 | 0.14 | 0.05 | 0.02 | 0.57 | 0.33 | 0.38 | – | 0.06 |
| SPL | 0.61 | 0.87 | 0.38 | 0.98 | 0.86 | 0.60 | 0.95 | 0.82 | 0.71 | 0.78 | – |

**Table S3:** Marker genotypes for the subset of 26 accessions from the Tusón population at the twelve significant flowering time loci detected in [Wisser et al., 2019]. Numeric codes 1, −1, and 0 for each marker genotypes indicates homozygous for allele1, allele2, and heterozygous genotype, respectively. Markers are as follows: M1 = PZE.101091520; M2 = PZE.101162348; M3 = PZE.102146567; M4 = PZE.104005434; M5 = PZE.104069513; M6 = PZE.105112537; M7 = SYN38278; M8 = SYN29958; M9 = PZE.108080696; M10 = ZmCCT10 CACTA; M11 = PZE.110050177; M12 = PZE.110055163. Acc = accession; gen = generation of selection; prop = proportion; het = heterozygous.

| Acc | Gen | M1 | M2 | M3 | M4 | M5 | M6 | M7 | M8 | M9 | M10 | M11 | M12 |
| --- | --- | --- | --- | --- | --- | --- | --- | --- | --- | --- | --- | --- | --- |
| 165.1 | 0 | -1 | . | 0 | 0 | -1 | 1 | 0 | 0 | 1 | 1 | 1 | 1 |
| 165.2 | 0 | -1 | 1 | -1 | 0 | . | 1 | 1 | 0 | 1 | 1 | 0 | 1 |
| 169.1 | 0 | -1 | 1 | -1 | -1 | 1 | 1 | 1 | 0 | 1 | 1 | 1 | 1 |
| 172.2 | 2 | -1 | 0 | -1 | 0 | . | 1 | 1 | -1 | 1 | 1 | 1 | 1 |
| 172.3 | 2 | 0 | 1 | -1 | 0 | 1 | 1 | 1 | -1 | 1 | 1 | 1 | 1 |
| 176.4 | 4 | -1 | 1 | -1 | 0 | 1 | 1 | 1 | 0 | 1 | 1 | 1 | 1 |
| 176.5 | 4 | -1 | 1 | -1 | 1 | . | 1 | 1 | 0 | 1 | 1 | 1 | 1 |
| 177.6 | 4 | -1 | 1 | -1 | 0 | 1 | 1 | 1 | 0 | 1 | 1 | 1 | 1 |
| 178.6 | 4 | -1 | . | -1 | 0 | . | 1 | 1 | 0 | 1 | 1 | 1 | 1 |
| 830.2 | 2 | -1 | 1 | 0 | 0 | 1 | 1 | 1 | 0 | 1 | 1 | 1 | 1 |
| 832.2 | 2 | -1 | 1 | -1 | 0 | . | 1 | 1 | 0 | 1 | 1 | 1 | 1 |
| 832.6 | 2 | 0 | 0 | -1 | 0 | . | 1 | 1 | -1 | 1 | 1 | 1 | 1 |
| 839.2 | 4 | -1 | 1 | -1 | 1 | 1 | 1 | 1 | 0 | 1 | 1 | 1 | 1 |
| 849.4 | 6 | -1 | 1 | -1 | 0 | 1 | 1 | 1 | 0 | 1 | 1 | 1 | 1 |
| 852.3 | 6 | -1 | 1 | -1 | 0 | 1 | 1 | 1 | 0 | 1 | 1 | 1 | 1 |
| 856.5 | 6 | -1 | 1 | -1 | 1 | . | 1 | 1 | -1 | 1 | 1 | 1 | 1 |
| 857.8 | 6 | 1 | 1 | 0 | 0 | 1 | 1 | 1 | -1 | 1 | 1 | 1 | 1 |
| 858.4 | 6 | 0 | -1 | -1 | 0 | 1 | 1 | 1 | 1 | 1 | 1 | 1 | 1 |
| 860.1 | 8 | -1 | 1 | -1 | 0 | 1 | 1 | 1 | 0 | 1 | 1 | 1 | 1 |
| 861.1 | 8 | -1 | 1 | -1 | 0 | 1 | 1 | 1 | 1 | 1 | 1 | 1 | 1 |
| 864.3 | 8 | -1 | 1 | -1 | 0 | 1 | 1 | 1 | 1 | 1 | 1 | 1 | 1 |
| 869.6 | 10 | 0 | 1 | -1 | 0 | 1 | 1 | 1 | -1 | 1 | 1 | 1 | 1 |
| 872.2 | 10 | -1 | 1 | -1 | 0 | 1 | 1 | 1 | 0 | 1 | 1 | 1 | 1 |
| 873.1 | 10 | -1 | 1 | -1 | 0 | 1 | 1 | 1 | 1 | 1 | 1 | 1 | 1 |
| count | Allele1 | 1 | 19 | 0 | 3 | 16 | 24 | 23 | 4 | 24 | 24 | 23 | 24 |
|  | Allele2 | 19 | 1 | 21 | 1 | 1 | 0 | 0 | 6 | 0 | 0 | 0 | 0 |
|  | Het | 4 | 2 | 3 | 20 | 0 | 0 | 1 | 14 | 0 | 0 | 1 | 0 |
| prop | Allele1 | 0.04 | 0.79 | 0.00 | 0.13 | 0.67 | 1.00 | 0.96 | 0.17 | 1.00 | 1.00 | 0.96 | 1.00 |
|  | Allele2 | 0.79 | 0.04 | 0.88 | 0.04 | 0.04 | 0.00 | 0.00 | 0.25 | 0.00 | 0.00 | 0.00 | 0.00 |
|  | Het | 0.17 | 0.08 | 0.13 | 0.83 | 0.00 | 0.00 | 0.04 | 0.58 | 0.00 | 0.00 | 0.04 | 0.00 |

**Figure S1:**
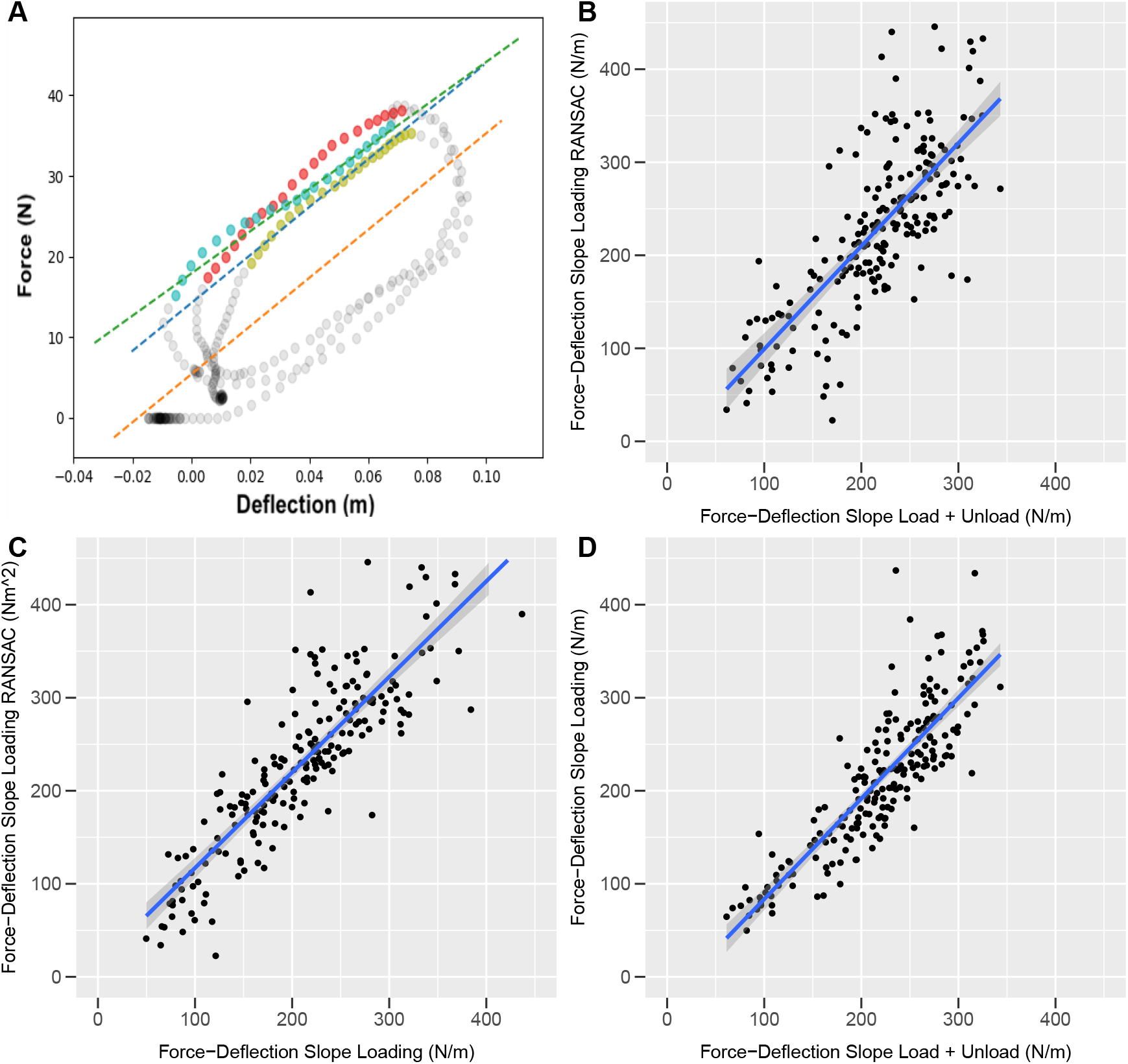
Comparison of Slope Extraction Methods. (A) Force-Deflection data was plotted for each test. In this representative image, the three cycles (different colors) that have been identified as loading due to the monotonically increasing deflections are shown by solid dots and grey dots represent unloading data. Dashed lines represent the three different slope fitting approaches: blue: ransac to the longest consecutive loading curve; green: line fit to all three loading curves; orange: line fit to loading and unloading data. (B) Correlation between the Force-Deflection slopes extracted by ransac fit to the loading data only and a line fit to loading and unloading data (r = 0.86, p < 2.2E^−16^). (C) Correlation between the Force-Deflection slopes extracted with the two loading methods (r = 0.87, p < 2.2E^16^). (D) Correlation between the Force-Deflection slopes extracted by a line fit to the loading data only and a line fit to loading and unloading data (r = 0.76, p < 2.2E^−16^). Lines in B, C and D represent generalized linear model (glm) fit and shading indicates a 95% confidence interval.

**Figure S2:**
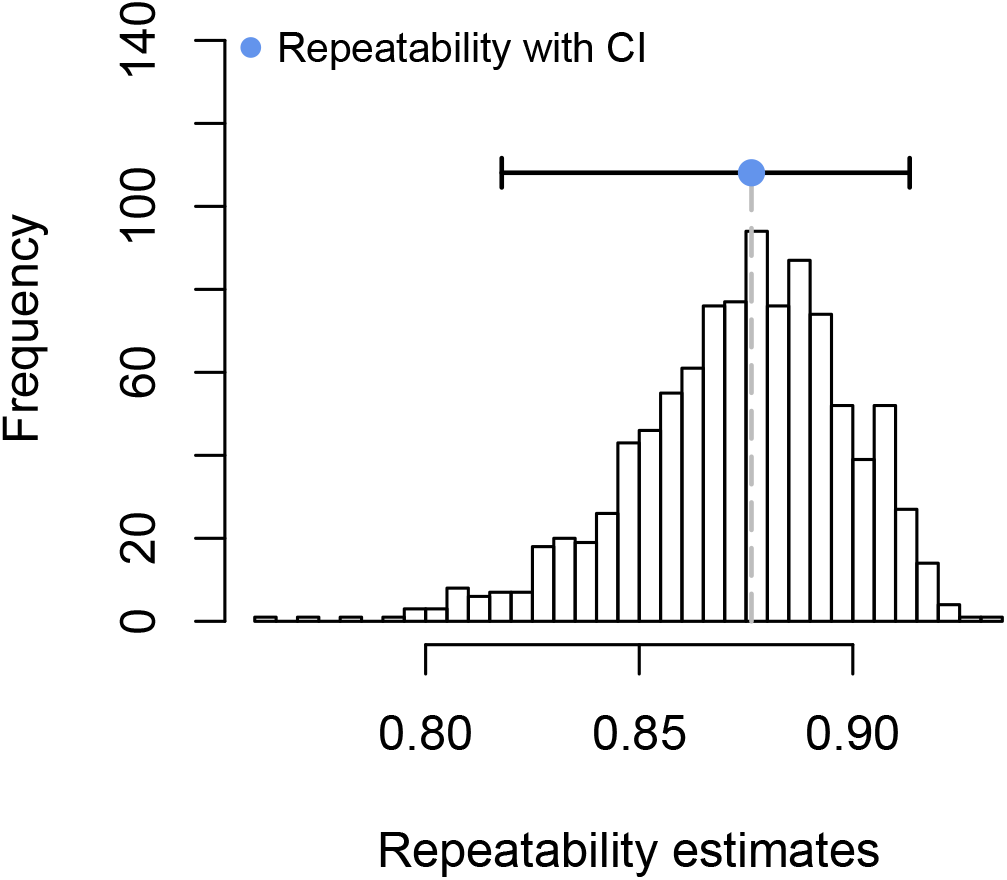
Measurements of the Force-Deflection Slope have High Repeatability. Repeatability analysis for plants in Figure 2 were calculated using the rptR package in R and confidence intervals determined by bootstrapping (n = 1000). The mean repeatability (R) was 0.876 with a 0.025 SE and p = 2.31E^−49^.

**Figure S3:**
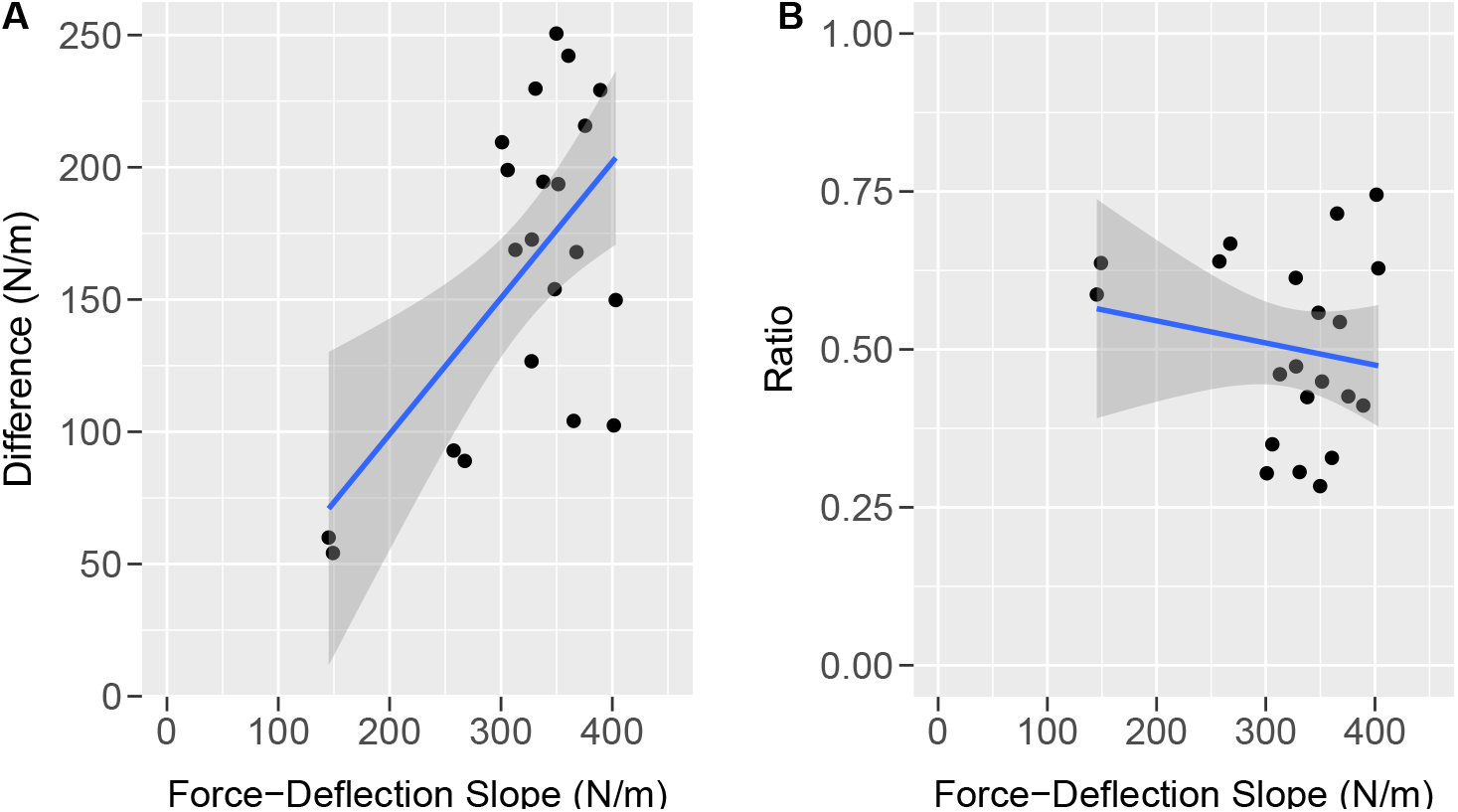
The Initial Force-Deflection Slope is Correlated with the Absolute, but Not the Relative Contribution of Brace Roots. (A) Comparison of the Force-Deflection slope with brace roots intact and the difference in Force-Deflection slope after all brace roots have been removed. There is a positive correlation (r = 0.59, p = 0.004) between these measurements. As the difference relies on the initial Force-Deflection slope, this correlation is to be expected. (B) Comparison of the Force-Deflection slope with brace roots intact and the ratio of Force-Deflection slope after all brace roots have been removed. There is no correlation (r = −0.17, p = 0.461) between these measurements. Lines represent generalized linear model (glm) fit and shading indicates a 95% confidence interval.

**Figure S4:**
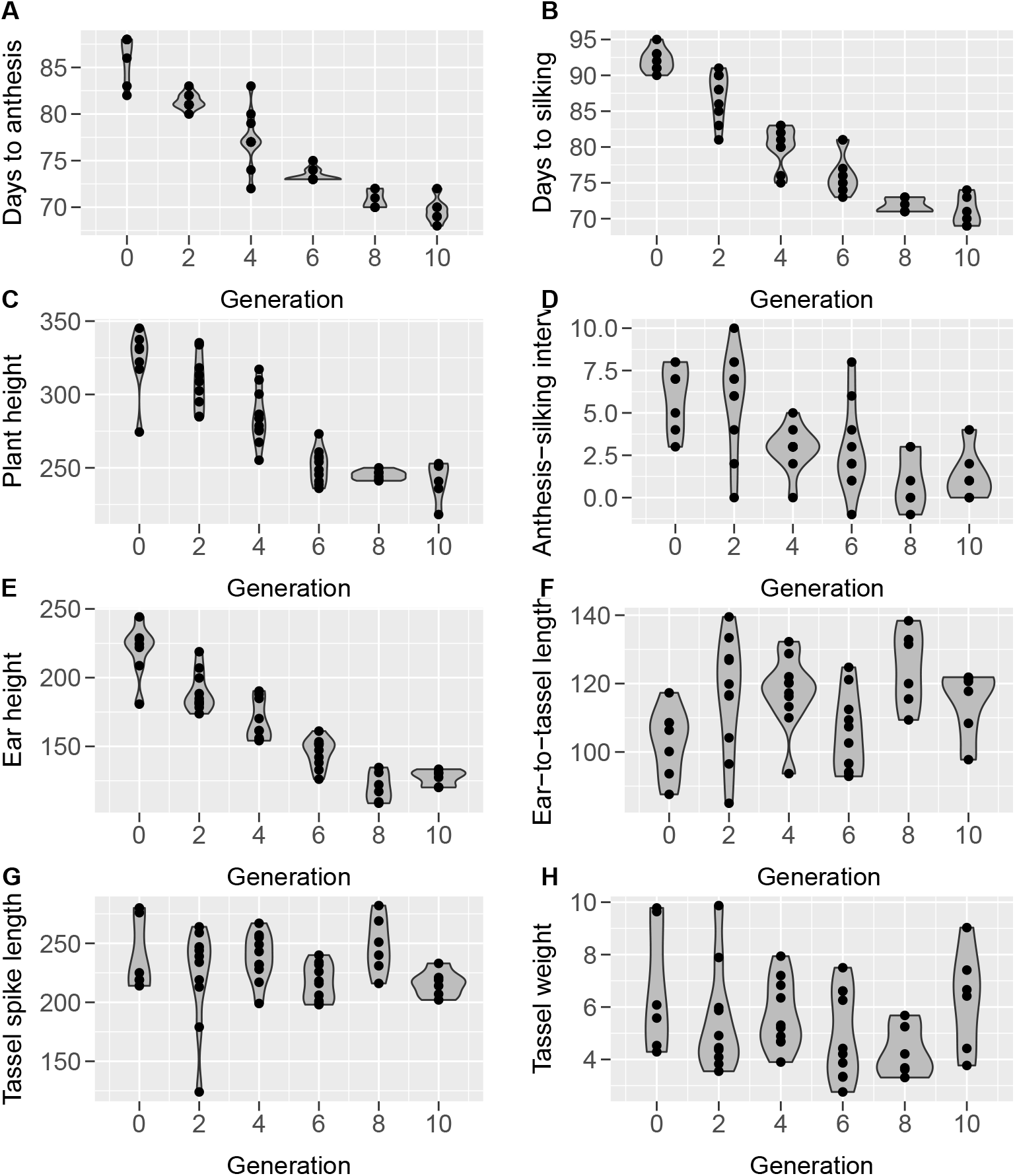
The subset of Tusón lines selected for this study show a reduction (A) days to anthesis, (B) days to silking, (B) plant height, (D) anthesis-silking interval, and (E) ear height. There was no change in (F) ear-to-tassel length, (G) tassel spike length, or (H) tassel dry weight in this subset of lines. Data from the Delaware grow out of these lines as published in Teixeira et al. [2015].

